# The seven transmembrane domain protein MoRgs7 functions in surface perception and undergoes coronin MoCrn1-dependent endocytosis in complex with Gα subunit MoMagA to promote cAMP signaling and appressorium formation in *Magnaporthe oryzae*

**DOI:** 10.1101/435511

**Authors:** Xiao Li, Kaili Zhong, Ziyi Yin, Jiexiong Hu, Lianwei Li, Haifeng Zhang, Xiaobo Zheng, Ping Wang, Zhengguang Zhang

## Abstract

Regulator of G-protein signaling (RGS) proteins primarily function as GTPase-accelerating proteins (GAPs) to promote GTP hydrolysis of Gα subunits, thereby regulating G-protein mediated signaling. RGS proteins could also contain additional domains such as GoLoco to inhibit GDP dissociation. The rice blast fungus *Magnaporthe oryzae* encodes eight RGS and RGS-like proteins (MoRgs1 to MoRgs8) that have shared and distinct functions in growth, appressorium formation and pathogenicity. Interestingly, MoRgs7 and MoRgs8 contain a C-terminal seven-transmembrane domain (7-TM) motif typical of G-protein coupled receptor (GPCR) proteins, in addition to the conserved RGS domain. We found that MoRgs7, together with Gα MoMagA but not MoRgs8, undergoes endocytic transport from the plasma membrane to the endosome upon sensing of surface hydrophobicity. We also found that MoRgs7 can interact with hydrophobic surfaces via a hydrophobic interaction, leading to the perception of environmental hydrophobic cues. Moreover, we found that MoRgs7-MoMagA endocytosis is regulated by actin patch-associated protein MoCrn1, linking it to cAMP signaling. Our studies provided evidence suggesting that MoRgs7 could also function in a GPCR-like manner to sense environmental signals and it, together with additional proteins of diverse functions, promotes cAMP signaling required for developmental processes underlying appressorium function and pathogenicity.

**Author summary:** The 7-TM domain is considered the hallmark of GPCR proteins, which activate G proteins upon ligand binding and undergo endocytosis for regeneration or recycling. Among eight RGS and RGS-like proteins of *M. oryzae*, MoRgs7 and MoRgs8 contain the 7-TM domain in addition to the RGS domain. We found that MoRgs7 can form hydrophobic interactions with the hydrophobic surface. This interaction is important in sensing hydrophobic cues by the fungus. We also found that, in response to surface hydrophobicity, MoRgs7 couples with Gα subunit MoMagA to undergo endocytosis leading to the activation of cAMP signaling. Moreover, we found that such an endocytic event requires functions of the actin-binding protein MoCrn1. Our results revealed MoRgs7 functions as a GPCR-like receptor protein to sense surface cues and activate signaling required for pathogenesis, providing new insights into G-protein regulatory mechanisms in this and other pathogenic fungi.

## Introduction

In the rice blast fungus *Magnaporthe oryzae*, the appressorium is a special infection structure produced by this fungus to penetrate the host plant. Appressorium formation and function depend on signal transduction pathway including G protein-coupled receptors (GPCRs)/G protein-mediated cAMP signaling [1, 2]. Once extracellular surface cues are sensed by GPCRs, such as the non-canonical GPCR Pth11 at the plasma membrane, the GPCR stimulates the specific G-protein Gα subunit for activating the cAMP signaling pathway [2]. *M. oryzae* contains three distinct Gα subunits (MoMagA, MoMagB, and MoMagC) [3, 4] and other conserved pathway components, such as adenylate cyclase MoMac1, cAMP-dependent protein kinase A catalytic subunits MoCpkA, and MoCpk2 [1, 5–7]. Together, they regulate not only growth but also appressorium formation and pathogenesis.

In addition, *M. oryzae* contain at least eight RGS (regulator of G-protein signaling) and RGS-like proteins (MoRgs1 to MoRgs8). Previous studies found that all these RGS proteins have certain regulatory functions in various aspects of growth and pathogenicity with MoRgs1, MoRgs2, MoRgs3, MoRgs4, MoRgs6, and MoRgs7 being mainly involved in appressorium formation and MoRgs1, MoRgs3, MoRgs4, and MoRgs7 in full virulence [3, 8]. Despite such understandings, detailed mechanisms associated with specific RGS proteins remain not understood, in particular, RGS-like MoRgs7 and MoRgs8 proteins that also contain a seven-transmembrane domain (7-TM). Previous studies have found that RGS proteins such as human Rgs14 contains a C-terminal GoLoco/G protein regulatory motif exhibiting an in vitro GDP-dissociation inhibitor for Gα(i) [9]. Since the 7-TM domain is a hallmark of GPCRs important in signal perception and transduction, we were interested in characterizing whether MoRgs7 or MoRgs8 has additional functions mimicking a GPCR. We found that MoRgs7, but not MoRgs8, is involved in a distinct regulating mechanism. MoRgs7 couples with MoMagA to undergo the endocytic process that is triggered by sensing surface hydrophobicity. Interestingly, MoRgs7 can interact with the hydrophobic surface to sense environmental hydrophobic cues. In addition, MoRgs7 endocytosis involves the actin-binding coronin homologue protein MoCrn1. Together, they contribute to G-protein/cAMP signaling required for appressorium function and pathogenicity.

## Results

### MoRgs7 function requires the RGS and 7-TM domains

Despite containing a relatively conserved RGS/RGS-like domain, MoRgs1-MoRgs8 of the blast fungus are structurally divergent [8]. MoRgs7 and MoRgs8, in particular, contain a long C-terminus domain that was analyzed by transmembrane domain prediction systems (http://mendel.imp.univie.ac.at/sat/DAS/DAS and www.cbs.dtu.dk/services/TMHMM) to have a GPCR-like 7-TM motif (S1 Fig and S2A Fig). MoRgs7 was demonstrated to have a role in appressorium function and pathogenicity, and this role is dependent on the 7-TM domain [8, 10].

To dissect the roles of MoRgs7 domains, the RGS domain was deleted (S2A Fig) and the mutant allele containing the 7-TM was fused to GFP and expressed in the Δ*Morgs7* mutant. The fusion proteins MoRgs7^Δ7-TM^:GFP and MoRgs7:GFP [10] were also expressed in the Δ*Morgs7* mutant as a control. Analysis results showed that the Δ*Morgs7* mutant expressing 7-TM:GFP still remained a relatively high cAMP concentration, similar to the Δ*Morgs7* mutant [8] but not the wild-type strain (S2B Fig). In hydrophobic surfaces, about 8% of Δ*Morgs7* conidia improperly generated two appressoria (S1C Fig), which could also be observed in the Δ*Morgs7*/*7-TM* strain (S2C Fig). The Δ*Morgs7*/*7-TM* strain was also attenuated in virulence, similar to the Δ*Morgs7* mutant (S2D and S2E Fig). In contrast, the expression of MoRgs7-GFP was able to suppress most of the defects in the Δ*Morgs7* strain. These tests showed that the 7-TM and RGS domains are important for MoRgs7 function in cAMP and virulence. However, the test failed to establish an independent role of the 7-TM.

### MoRgs7 and Gα protein MoMagA undergo internalization in response to the exposure to hydrophobic surfaces

MoMagA plays a major role in cAMP signaling, appressorium formation and pathogenesis in *M. oryzae* and it is also one of the three Gα subunits demonstrated to interact with MoRgs7 [8]. To investigate additional functional mechanisms of MoRgs7-MoMagA interaction, we first validated the interaction through co-immunoprecipitation (co-IP). In addition to MoMagA, the constitutively activated MoMagA^G187S^ allele [3] was also included in the test. The result showed that MoRgs7 can interact with both MoMagA and MoMagA^G187S^ alleles and that the 7-TM and the RGS domain both can interact with MoMagA (Fig 1A and 1B).

**Fig 1.**
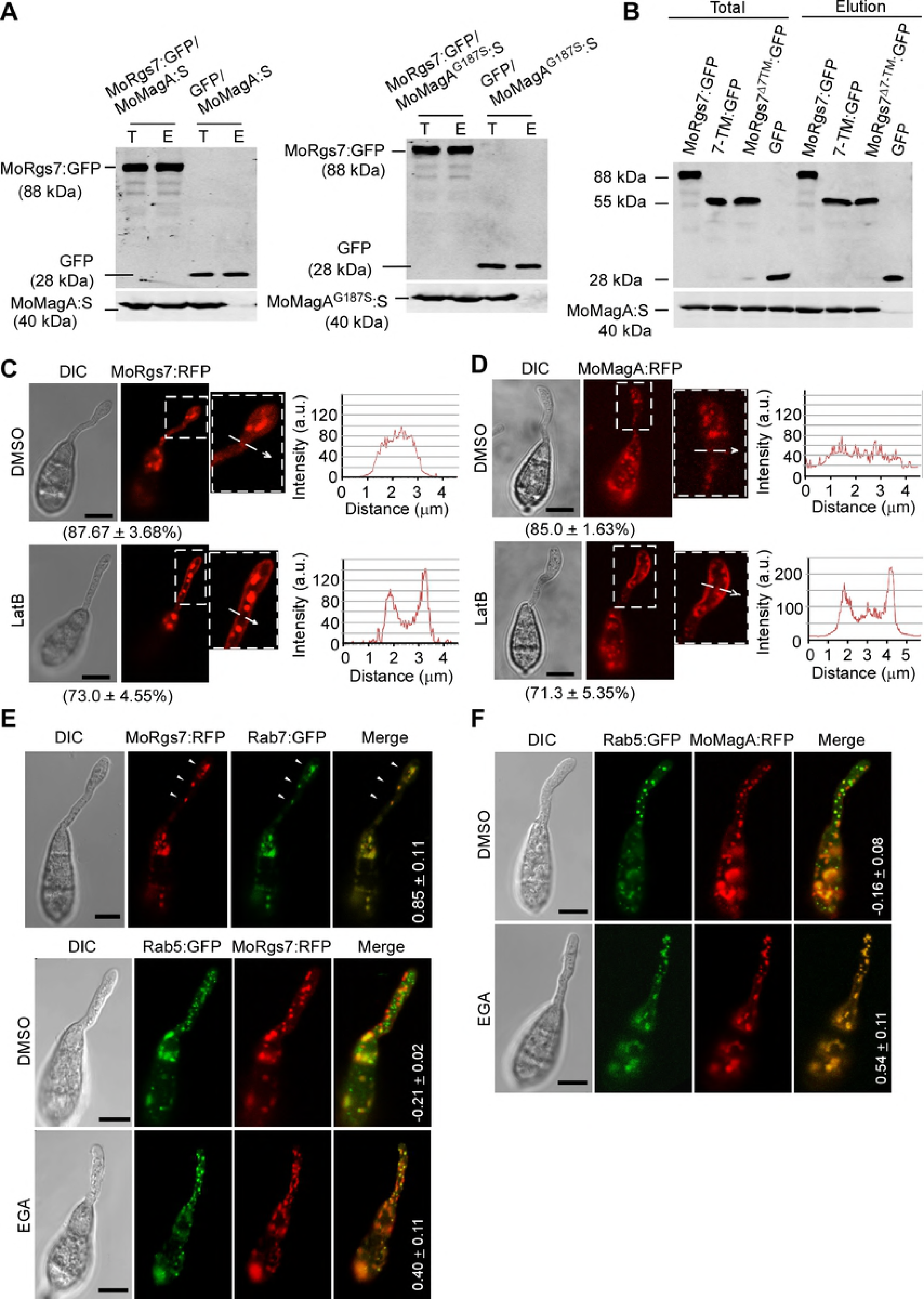
Endocytosis internalizes MoRgs7 and MoMagA from the plasma membrane to endosomes. **(A)**Co-IP assay for examining the interaction between MoRgs7 and MoMagA or MoMagA^G187S^. T represents total proteins, which were extracted from the mycelium of the strain expressing MoRgs7:GFP or GFP and MoMagA:S or MoMagA^G187S^:S. Total proteins were used for incubation with anti-GFP beads. E represents elution, which contains the proteins precipitated with GFP fusion proteins. GFP protein was used as negative control. These samples were probed using anti-GFP and anti-S antibodies. **(B)**Co-IP assay for examining the interaction of MoRgs7, 7TM of MoRgs7 and MoRgs7^Δ7TM^ with MoMagA. The total proteins were extracted from the mycelium of the strains co-expressing MoRgs7:GFP and MoMagA:S, 7TM:GFP and MoMagA:S, and MoRgs7^Δ7TM^:GFP and MoMagA:S, respectively. The elution contains the proteins precipitated with GFP fusion proteins.. These samples were probed using anti-GFP and anti-S antibodies. **(C and D)**LatB enhanced MoRgs7:RFP and MoMagA signals at the plasma membrane of germ tubes (3 h post-inoculation). The DMSO solvent treatment was used as a control. The sections selected by white arrows were subjected to linescan analysis for the distributions of MoRgs7:RFP and MoMagA:RFP. Percentage of a pattern showed in image was calculated by observation for 50 germinated conidia that were randomly chosen, and observation was conducted for 3 times. Bars = 5 μm. **(E and F)**EGA induced the localizations of MoRgs7:RFP and MoMagA:RFP on MoRab5-labeled early endosomes (3 h post-inoculation). The colocalization of MoRgs7:RFP with MoRab7:GFP and DMSO treatment were used as the control. The values were the mean Pearson’s correlation coefficients of MoRgs7:RFP/ MoMagA:RFP with MoRab5:GFP/MoRab7:GFP. ImageJ software was used to calculate the Pearson’s correlation coefficients, which were generated by analyzing images taken from 5 germinated conidia. Bars = 5 μm.

Since GPCRs undergo endocytosis for receptor recycling [11], and both of MoRgs7 and MoMagA were localized to late endosomes that are the main components of the endocytic pathway, we hypothesized that MoRgs7 and MoMagA may also undergo actin-dependent endocytosis. To test this, we employed actin polymerization inhibitor latrunculin B (LatB) to disrupt endocytosis as previously described [12, 13]. At 3 h post-inoculation, MoRgs7:RFP and MoMagA:RFP signals remained very strong at the plasma membrane (PM) of the germ tube, in contrast to DMSO control (Fig 1C and 1D). Given that 4-bromobenzaldehyde *N*-2,6-dimethylphenyl (EGA) inhibits early to late endosome transport [14], it was applied that led to an appearance of MoRgs7 and MoMagA RFP signals in MoRab5:GFP-labeled early endosomes in germ tubes, in contrast to DMSO control (Fig 1E and 1F). Without EGA treatment, MoRgs7:RFP was predominantly localized to Rab7:GFP-labeled late endosomes (Fig 1E). These co-localizations of proteins with endosomes were corroborated by Pearson correlation coefficient statistical analysis. Taken together, MoRgs7 and MoMagA movement follows the common endocytic pathway.

To further validate MoRgs7 and MoMagA endocytosis, we photobleached the MoRgs7 and MoMagA fluorescence in late endosomes of the germ tubes on hydrophobic surfaces and examined the fluorescence recovery dynamic using Fluorescence Recovery After Photobleaching (FRAP) at 3 h post-inoculation. In addition, we applied the microtubule-destabilizing benomyl to inhibit endosome trafficking via microtubule and cycloheximide to inhibit newly synthesized fluorescent proteins moving into endosomes [15]. We found that endocytosis promotes recovery of RFP fluorescence of MoRgs7 and MoMagA in late endosomes within 90 sec (Fig 2A and 2B). Furthermore, we used FRAP to bleach the fluorescence in endosomes in the germ tube on the hydrophilic surfaces at 3 h post-inoculation. The recovery of fluorescence of MoRgs7:RFP and MoMagA:RFP in the endosomes was rarely detected (Fig 2A and 2B), suggesting that MoRgs7 and MoMagA are rarely internalized through endocytosis upon the perception of the hydrophilic surface.

**Fig 2.**
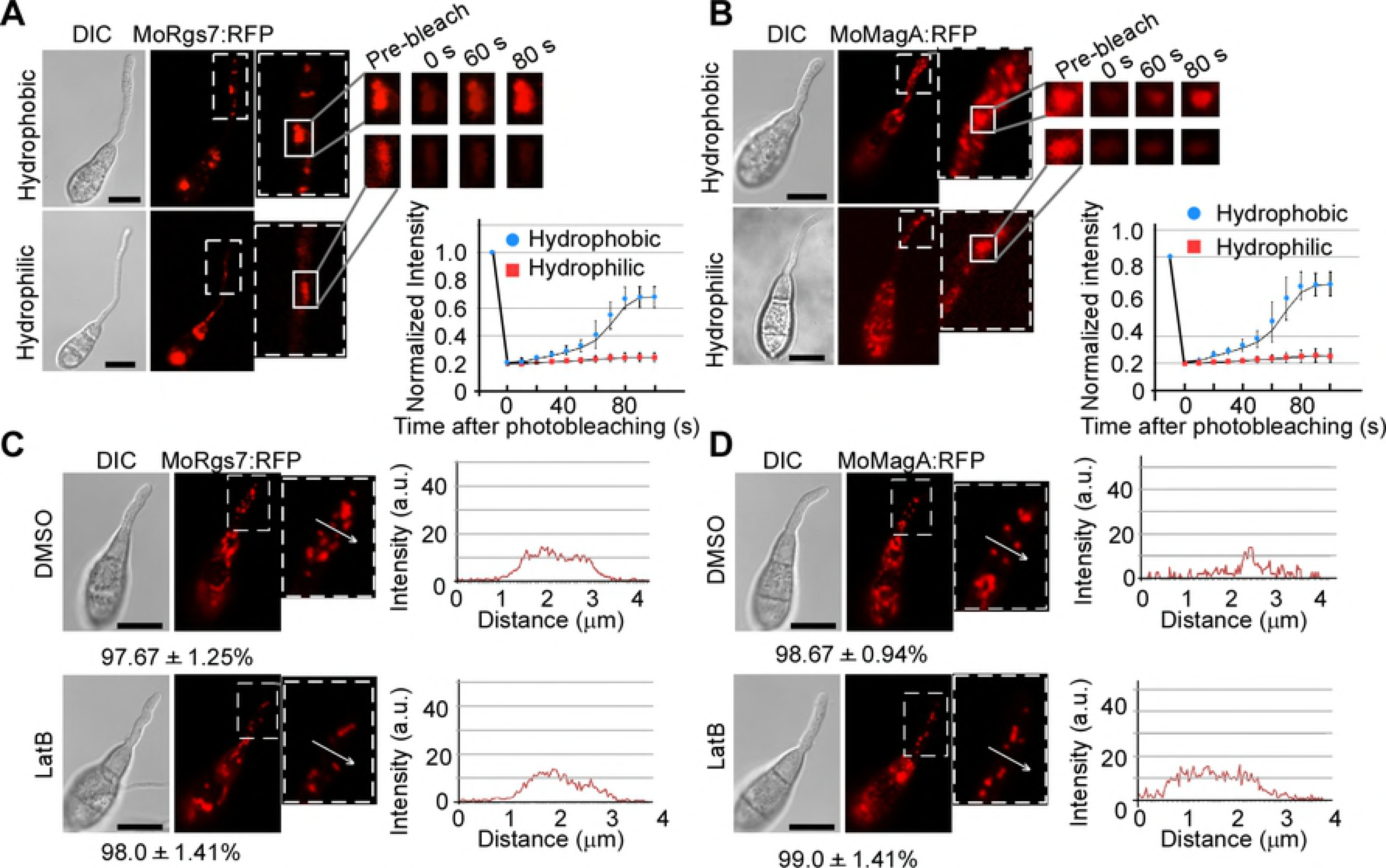
MoRgs7 and MoMagA endocytosis is active when *M. oryzae* germinates on hydrophobic surfaces, not hydrophilic surfaces. **(A and B)** FRAP for measuring the recovery of MoRgs7:RFP and MoMagA:RFP on endosomes at 3 h-post inoculation. The conidia were allowed to germinate on hydrophobic and hydrophilic surfaces. The representative images of FRAP were shown and the selected areas were measured for fluorescence recovery after photobleaching. The normalized FRAP curve was fitted with measuring 15 regions from different cells. Bars = 10 μm. **(C and D)** For the germinated conidia on hydrophilic surfaces, LatB could not induce the accumulation of MoRgs7:RFP and MoMagA:RFP at the plasma membrane of germ tubes at 3 h-post inoculation. The sections selected by white arrows were subjected to linescan analysis for the distributions of MoRgs7:RFP and MoMagA:RFP. Percentage of a pattern showed in image was calculated by observation for 50 germinated conidia that were randomly chosen, and observation was conducted for 3 times. Bars = 10 μm.

Intriguingly, the absence of MoRgs7 and MoMagA endocytosis on the hydrophilic surface did not couple with accumulation of MoRgs7:RFP or MoMagA:RFP signals at the plasma membrane (PM) of the germ tubes (Fig 2C and 2D). As treating germinated conidia with LatB on hydrophilic surfaces for 1 h still could not cause accumulation of RFP signals at the PM (Fig 2C and 2D), we thus reasoned that in response to exposure to hydrophilic cues MoRgs7 and MoMagA were rarely sent to the PM from intracellular systems.

### MoRgs7 localization pattern is different from MoRgs8

MoRgs8 also contains 7-TM domain. To examine whether MoRgs8 undergoes similar endocytosis, we expressed MoRgs8:GFP in Guy11 and observed MoRgs8 localization during appressorium development on the hydrophobic surface. However, MoRgs8:GFP was found evenly distributed in the cytoplasm of germ tubes (Fig 3A). When compared with MoRgs7:GFP (Fig 3B), MoRgs8:GFP did not display any obvious endosome-localization patterns in the germ tubes. Further, LatB failed to cause any effects to MoRgs8:GFP distribution (Fig 3A). In contrast, the MoRgs7:GFP signal was enhanced at the PM in response to LatB (Fig 3B). These results revealed that MoRgs8 may function differently from MoRgs7.

**Fig 3.**
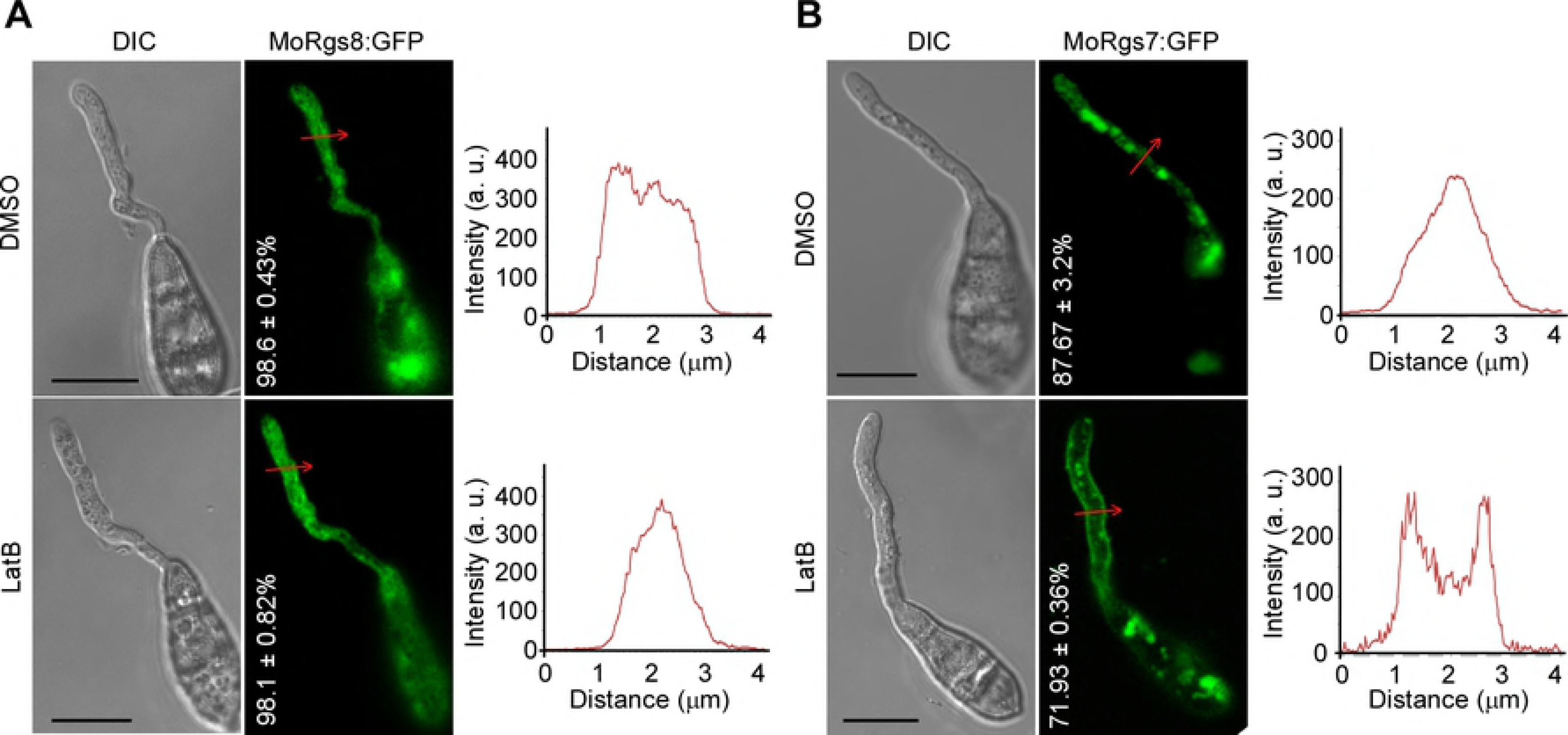
MoRgs8 does not undergo endocytosis. **(A)** LatB could not induce the accumulation of MoRgs8:GFP at the plasma membrane of germ tubes on the hydrophobic surface at 3 h-post inoculation. Percentage of a pattern showed in image was calculated by observation for 50 germinated conidia that were randomly chosen, and observation was conducted for 3 times. Bars = 10 μm. **(B)** LatB enhanced the signals of MoRgs8:GFP at the plasma membrane of germ tubes on hydrophobic surface at 3 h-post inoculation, which was the control of the treatment to MoRgs8:GFP. Bars = 10 μm.

### MoRgs7 is able to interact with the hydrophobic surface

Since the above results showed that the hydrophobic surface, not the hydrophilic surface, induces the PM localization of MoRgs7 in germ tubes during appressorium development, we hypothesized that MoRgs7 is possibly involved in sensing hydrophobic surfaces and the 7-TM may have a role in this process. We hypothesized that MoRgs7 at the PM may attach to hydrophobic surfaces in a hydrophobic interactive manner, and formation of such interactions by PM proteins including MoRgs7 is a step in the perception of hydrophobic cues.

To test this hypothesis, we first examined whether MoRgs7 has the ability to bind to hydrophobic materials by performing an affinity precipitation assay with phenyl-agarose gel beads. The phenyl groups attached to the beads are highly hydrophobic. The beads were then incubated with MoRgs7:GFP and the GFP protein (a negative control), respectively, in a high concentration of salt solution containing 1.5 M NaCl and 1.5 M MgSO_4_. This allowed proteins to bind to the beads, due to that at high salt concentration of non-polar side chains on the surface upon protein interactions with the hydrophobic groups [16]. Then we washed the beads to remove unbound proteins using a series of aqueous solutions with different salt concentration. If an intense hydrophobic interaction between the protein and phenyl groups was formed, the protein will be hardly removable from beads even by low salt concentration solution containing 0.3 M NaCl and 0.3 M MgSO_4_, or containing 0.2 M NaCl and 0.2 M MgSO_4_. After washing, we used Western-blot analysis to detect the amount of MoRgs7:GFP or GFP that remained bound to beads. The results indicated that MoRgs7:GFP, but not GFP, remained in the elution (Fig 4A). This suggested that MoRgs7 has a strong ability to interact with hydrophobic materials and this ability may allow MoRgs7 to mediate a hydrophobic interaction between the pathogen and the hydrophobic surface.

**Fig 4.**
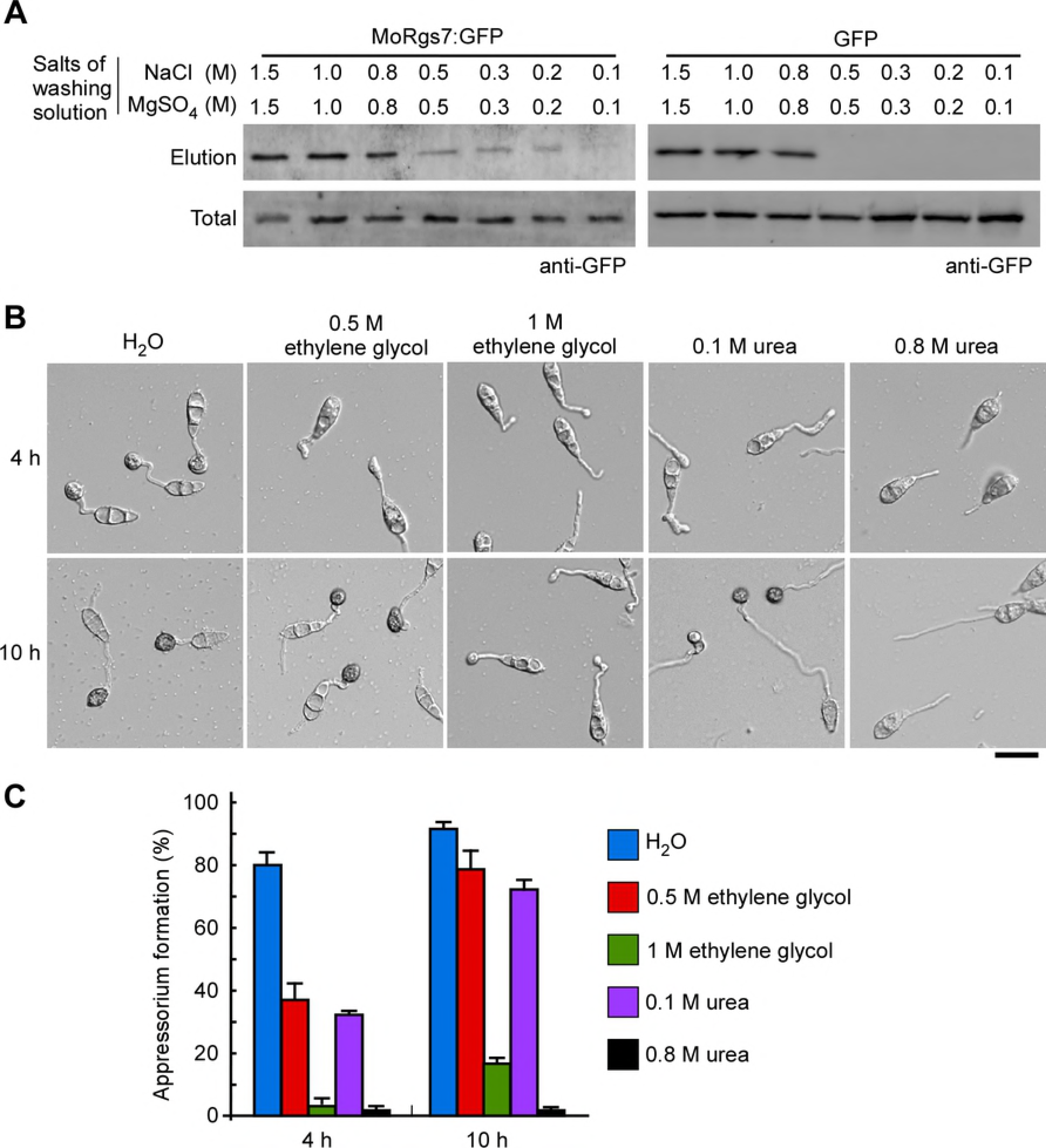
Formation of a hydrophobic interaction with hydrophobic surfaces is mediated by MoRgs7 that is important for the perception of hydrophobic cues. **(A)** Affinity precipitation experiments show that MoRgs7 intensely binds to hydrophobic groups that attach to beads. GFP proteins were used as control. A series of washing solutions with different salt concentrations were used to wash off the unbound proteins. The MoRgs7:GFP or GFP protein precipitated by beads was present in elution, which was detected by anti-GFP antibody. **(B)** Appressorium formation assay was performed with treatments of ethylene glycol and urea solutions. The images were taken at 4 h and 10 h post-inoculation. Bar = 20 μm. **(C)**The percentages of appressorium formation were calculated by observation for 100 germinating conidia that were randomly chosen and the observation was conducted for 3 times.

We then investigated whether MoRgs7 forming a hydrophobic interaction with hydrophobic surfaces is an approach of *M. oryzae* to detect hydrophobic cues. Given that urea and ethylene glycol can interrupt hydrophobic interactions by causing a disordering of water molecules on hydrophobic regions [17, 18], they were applied to germinating conidia on hydrophobic surfaces at 1 h-post inoculation when most of conidia only germinated with a germ tube. In the presence of 0.5 M ethylene glycol or 0.1 M urea, appressorium formation was about 50% lower than that of water treatment at 4 h post-inoculation, even though almost 80% of conidia developed appressorium at 10 h post-inoculation (Fig 4B and 4C). Moreover, in the presence of 1 M ethylene glycol or 0.8 M urea, less than 20% of conidia developed appressoria even at 10 h post-inoculation. Most of conidia only germinated germ tubes with or without swelling at terminals. These results implied that a successful hydrophobic interaction formation is a critical step in hydrophobic surface recognition by *M. oryzae*.

### MoRgs7 and MoMagA are independent of each other in endocytosis

To examine the nature of MoRgs7-MoMagA endocytosis and whether MoRgs7 internalization is dependent on MoMagA, we determined the rate of MoRgs7 internalization in the wild-type strain Guy11 and the Δ*MoMagA* mutant using FRAP analysis. We found that MoRgs7 internalization had a normal rate in the Δ*MoMagA* as that in Guy11 (Fig S3A and S3B). In addition, the internalization rate of MoMagA was also the same in Guy11 and the Δ*Morgs7* strain (Fig S3C and S3D). These results suggested that MoRgs7 and MoMagA internalizations were independent of each other.

### MoCrn1 interacts with MoRgs7 and F-actin, and affects microtubule function

To further understand the endocytosis process of MoRgs7, we searched for additional protein partners of MoRgs7 through a yeast two-hybrid (Y2H) screening and identified a coronin protein homolog, MoCrn1, as two polypeptides of 148 and 273 amino acids, from a cDNA library in the pGADT7 vector. MoRgs7 cDNA was inserted into pGBKT7 as bait. The interaction was specific, as an interaction between MoCrn1 and other RGS proteins, including MoRgs1, MoRgs3 and MoRgs4, cannot be established (Fig 5A). The interaction was further validated by co-IP and bimolecular fluorescence complementation (BiFC). The co-IP assay indicated that both the 7-TM and the RGS domains could interact with MoCrn1, independently (Fig 5B and 5C). BiFC revealed that MoCrn1 interacts with MoRgs7 during appressorium development (Fig 5D). The YFP signal could be detected at the PM while some weak signals appeared in the cytoplasm (Fig 5D).

**Fig 5.**
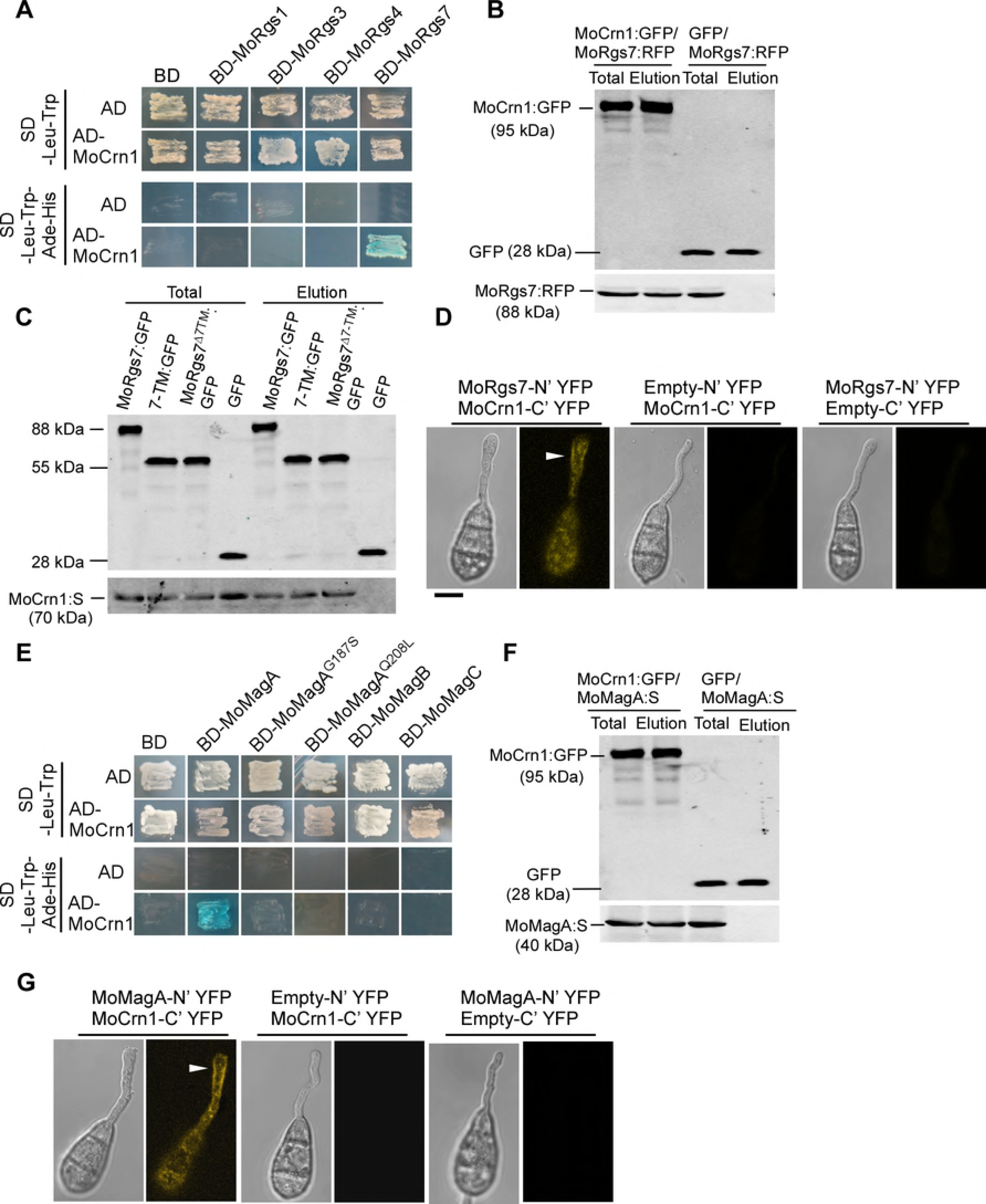
MoCrn1 interacts with MoRgs7 and MoMagA. **(A)** The yeast two-hybrid assay for examining the interaction of MoCrn1 with MoRgs1, MoRgs3, MoRgs4, and MoRgs7. The yeast transformants were isolated from SD-Leu-Trp plates, following growing on SD-Leu-Trp-His-Ade plates containing X-α-Gal for examining β-galactosidase activity. The transformants expressing AD and BD, BD-MoRgs1/3/4/7 and AD, BD and AD-MoCrn1 were used as negative control. **(B)** Co-IP assay for examining the interaction between MoCrn1 (tagged with GFP) and MoRgs7 (tagged with RFP tag) using anti-GFP beads. The total proteins were extracted from the mycelium of the strain co-expressing MoCrn1-GFP and MoRgs7-RFP. GFP protein was used as negative control. The anti-GFP and anti-RFP antibodies were used to detect GFP, MoCrn1-GFP and MoRgs7-RFP proteins. **(C)** Co-IP assay for examining the interaction of MoRgs7, 7-TM and MoRgs7^Δ7TM^ (tagged with GFP) with MoCrn1 (tagged with S-tag) using anti-GFP beads. **(D)** BiFC assay for examining the interaction of MoCrn1 with MoRgs7. The conidia were allowed to germinate on hydrophobic surfaces for 3 h. YFP signals could be detected from the germ tubes and conidia of the strain co-expressing MoRgs7-N’YFP and MoCrn1-C’YFP. The strains co-expressing empty-N’YFP and MoCrn1-C’YFP, and co-expressing MoRgs7-N’YFP and Empty-C’YFP were used as negative control. Bar = 5 μm. **(E)** The yeast two-hybrid assay for examining the interaction of MoCrn1 with MoMagA, MoMagB, MoMagC and two activated forms of MoMagA, MoMagA^G187S^ and MoMagA^Q208L^. **(F)** Co-IP assay for examining the interaction of MoCrn1 (tagged with GFP) with MoMagA (tagged with S-tag) using anti-GFP beads. **(E)** BiFC assay for examining the interaction of MoCrn1 and MoRgs7. YFP signals could be detected from the germ tubes and conidia of the strain co-expressing MoMagA-N’YFP and MoCrn1-C’YFP. The strains co-expressing empty-N’YFP and MoCrn1-C’YFP, and co-expressing MoMagA-N’YFP and Empty-C’YFP were used as negative control. Bar = 5 μm. **(G)** Co-IP assay for examining the interaction of MoCrn1 (tagged with GFP) with MoMagA (tagged with S-tag) using anti-GFP beads.

In the eukaryotic cells, coronin proteins act as F-actin binding proteins and regulate actin-related processes such as membrane trafficking [19]. We tested whether MoCrn1 associates with actin in *M. oryzae* using Lifeact, a living cell actin marker described previously [13, 20, 21]. The MoCrn1:GFP and Lifeact:RFP were co-expressed in Guy11 and co-localization of MoCrn1:GFP and Lifeact:RFP was examined under a confocal microscope. We observed that MoCrn1:GFP and actin were dispersed in nascent appressoria after 6 h of incubation (S5A Fig), and that MoCrn1 punctate patches were localized to the membrane. However, MoCrn1:GFP formed ring-like structures in mature appressoria, which were highly co-localized with the F-actin network at the center of mature appressoria (S5A Fig). We also observed that MoCrn1 patches were co-localized with actin patches at the hyphal tips and conidia (S5A Fig). The interaction between MoCrn1 and F-actin was again demonstrated through Y2H (S5B and S5C Fig).

We next investigated whether MoCrn1 affects the actin organization by generating a Δ*Mocrn1* mutant, in which *MoCRN1* gene knock-out was validated by Southern-blot (S4 Fig), and expressing Lifeact:RFP in the Δ*Mocrn1* mutant and Guy11. In Guy11, the hyphal tip regions were occupied with many actin patches that are associated with the PM (S5D Fig). However, about 20% of the hyphae formed some abnormal, enlarged actin patches in the cytoplasm of Δ*Mocrn1* (S5D Fig). Also, the enlarged actin patches could be found in over 10% of Δ*Mocrn1* conidia (S5E Fig), likely due to actin aggregation. Moreover, we found that Guy11 formed normal ring-like actin structure at the base of 80% appressoria, compared to 72% in Δ*Mocrn1* that displayed a disorganized actin network. This observation was confirmed by line-scan analysis (S5F Fig). Thus, we concluded that MoCrn1 regulates actin assembly and *MoCRN1* deletion caused minor defects in actin structures.

In the budding yeast *Saccharomyces cerevisiae*, Crn1 interacts with the microtubule [22]. The Δ*crn1* mutant cells as well as cells overexpressing Crn1 showed microtubule defects and the mutant Δ*crn1* is more sensitive than wild type strains to benomyl [23]. To determine whether MoCrn1 also affects the microtubule, the pYES2 construct containing the full-length MoCrn1 cDNA was expressed in the yeast Δ*crn1* mutant. On SD plates containing 10, 20, and 30 μg/ml benomyl, Δ*crn1* exhibited most significant inhibition in growth compared to the wild type strain BY4741 (S5G Fig). However, there was no significant difference between the Δ*crn1* strain expressing *MoCRN1* and BY4741. Further, we examined Guy11, the Δ*Mocrn1* mutant, and the complemented strain for benomyl resistance. On CM plates with 10, 20 and 30 μg/ml benomyl, we found that Δ*Mocrn1* was less sensitive to benomyl than Guy11 and the complemented strain (S5H Fig). Together, these results suggested that MoCrn1 has conserved microtubule-related functions.

### MoCrn1 is important for the internalization of MoRgs7 and MoMagA during appressorium development

As MoCrn1 interacts with MoRgs7 and is localized to actin patches that represent endocytic pits [24], we hypothesized that MoCrn1 may function as an adaptor protein to direct MoRgs7 for internalization. Therefore, we tested whether MoCrn1 affects the endocytosis of MoRgs7 by observing the spatial distribution of MoRgs7:RFP in germinated conidia on the hydrophobic surface at 3 h post-inoculation. Despite of that endosome-localized MoRgs7 was found in both the Δ*Mocrn1* mutant and Guy11, the Δ*Mocrn1* mutant displayed a higher concentration of MoRgs7:RFP at the PM of the germ tube than Guy11 did (Fig 6A). FRAP analysis indicated the fluorescence recovery of MoRgs7:RFP in Δ*Mocrn1* was evidently delayed than that in Guy11 (Fig 6C), suggesting that the diffusion of MoRgs7:RFP fluorescence into endosomes was impaired. This is consistent with our hypothesis that MoCrn1 is implicated in MoRgs7 internalization during appressorium development.

**Fig 6.**
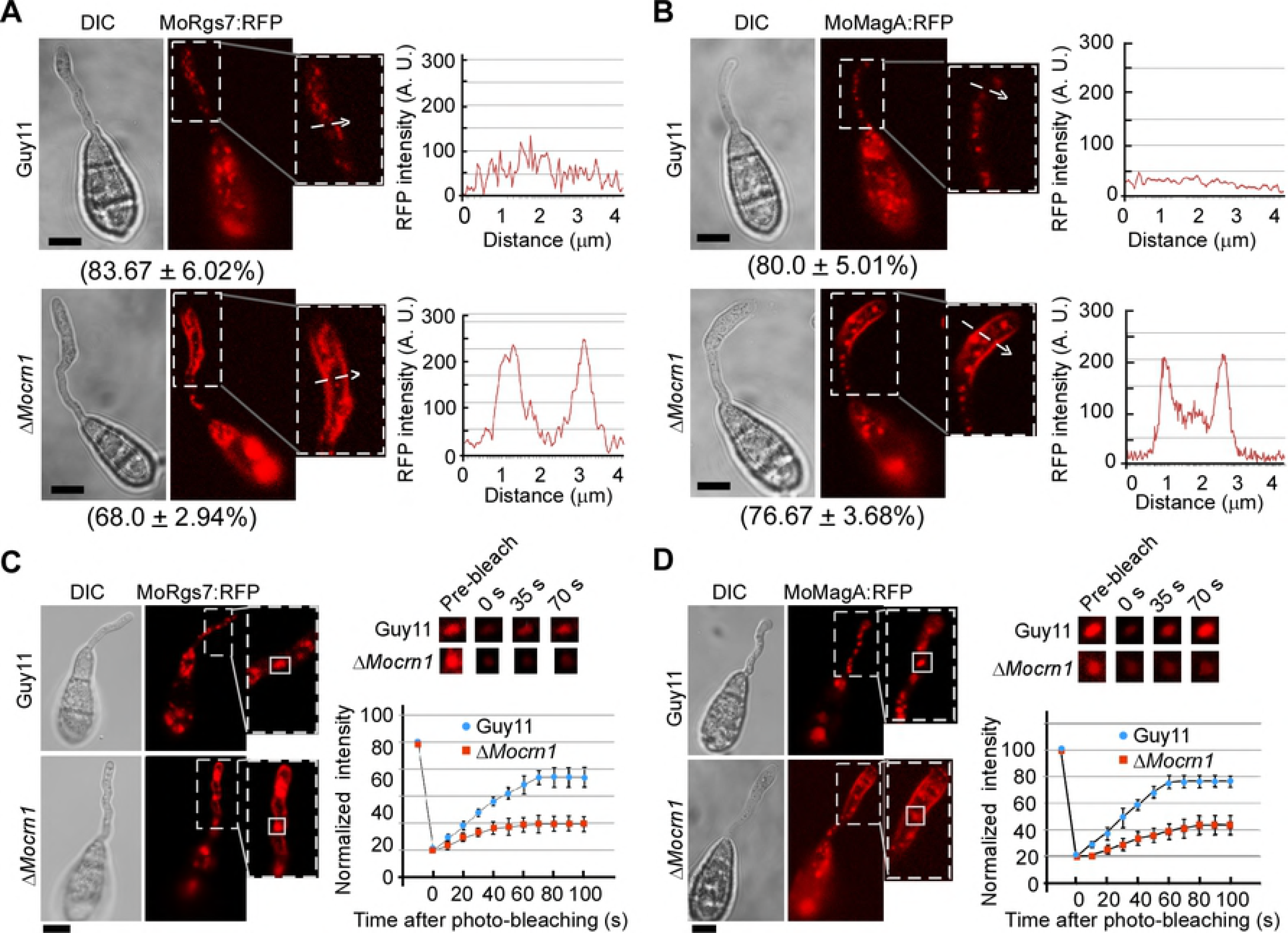
MoCrn1 regulates the endocytosis of MoRgs7 and MoMagA. **(A and B)** Images show the MoRgs7:RFP and MoMagA distributions in the germ tube of Guy11 and Δ*Mocrn1* at 3 h post-inoculation. White arrows indicated the regions in where the fluorescence intensity was measured by line-scan analysis. Percentage of a pattern showed in image was calculated by observation for 50 germinated conidia that were randomly chosen, and observation was conducted for 3 times. Bar = 5 μm. **(C and D)** FRAP for measuring the recovery of MoRgs7:RFP and MoMagA:RFP fluorescence in the endosomes of germ tubes. FRAP analysis was conducted at 3 h post-inoculation. The representative images of FRAP were shown and the selected areas were measured for fluorescence recovery after photobleaching. Bar = 5 μm. The normalized FRAP curve was fitted with measuring 15 regions from different cells.

Since MoRgs7 and MoMagA are both internalized via endocytosis, we also examined if MoCrn1 has a role in the MoMagA internalization through a protein-protein interaction. We first validated the interaction between MoCrn1 and MoMagA. In Y2H, we found that MoCrn1 interacts with MoMagA and this interaction was specific, since MoCrn1 was not found to interact with MoMagB and MoMagC (Fig 5E). In addition, MoCrn1 did not interact with MoMagA^G187S^ and MoMagA^Q208L^ (Fig 5E), the two-active forms of MoMagA [3]. The interaction between MoCrn1 and MoMagA was again confirmed by co-IP (Fig 5F) and BiFC assays (Fig 5G).

We next tested whether MoCrn1 affects the MoMagA distribution during appressorium development on hydrophobic surfaces. In Guy11, we have observed that MoMagA:RFP displayed the endosome localization pattern in germ tubes and conidia. In Δ*Mocrn1*, we could still observe MoMagA:RFP on late endosomes, but there was a significant increase in the membrane localization of MoMagA:RFP (Fig 6B). We again employed the FRAP assay to determine MoMagA internalization and found that the recovery of fluorescence of MoMagA:RFP in endosomes was slower in Δ*Mocrn1* than that in Guy11 (Fig 6D). These results confirmed that MoCrn1 is important for MoMagA internalization during appressorium development.

### MoCrn1 controls MoCap1 localization

MoCrn1 co-localizes with F-actin that is similar to the adenylate cyclase associated protein MoCap1 that functions in cAMP signaling [6]. To examine whether MoCrn1 is required for MoCap1 localization, we expressed MoCap1:GFP in Δ*Mocrn1* and observed that the actin-like localization pattern of MoCap1 was completely disrupted in appressoria, conidia and hyphae of Δ*Mocrn1* (S7 Fig). Strikingly, MoCap1 preferred to form cytoplasmic aggregations. Additionally, we found that MoCrn1 interacts with MoCap1 by performing a co-IP assay (Fig 8F), in which the strain co-expressing MoCrn1:GFP and MoCap1:S was used. These results led us to conclude that MoCrn1 has a crucial role in recruiting MoCap1 to actin patches.

### MoCrn1 is important for turgor generation, degradation of glycogen and lipid in the appressorium and pathogenicity

MoCrn1 has been associated with MoRgs7, MoMagA, and MoCap1 that all have a role in cAMP signaling. Indeed, we found that the Δ*Mocrn1* mutant also showed attenuated cAMP levels (S6A Fig) and a delay in appressorium formation (S8 Fig). At 4 h post-germination, nearly 40% of Δ*Mocrn1* conidia formed appressorium on a hydrophobic surface compared with 80% of Guy11 did. However, over 80% of Δ*Mocrn1* conidia could still form the appressorium at 6 h post-germination (S8 Fig). An incipient collapse assay indicated that MoCrn1 contributes to full turgor generation, since the collapse rate of the appressorium was significantly higher in Δ*Mocrn1* than in Guy11 and the complemented strains (S6B Fig).

Intracellular cAMP levels regulate the degradation of glycogen and lipid that are required for proper turgor generation in the appressorium [5, 25]. We thus compared the degradation of glycogen and lipid between the Δ*Mocrn1* mutant and Guy11 strains. Conidia were allowed to germinate on hydrophobic surfaces and iodine and Neil Red were used to stain glycogen and lipid, respectively [26]. At 6 h post-inoculation, glycogen appeared in the early appressorium (S6C Fig), and it broke down in 68.4% of the Guy11 appressoria after 16 h and 87% after 24 h, in comparison to 22.4% of Δ*Mocrn1* appressoria after 16 h and 53% after 24 h (S6E Fig). Resembling to the glycogen, lipid degradation in Δ*Mocrn1* appressoria was slower than Guy11. Lipid bodies disappeared in 44% of Δ*Mocrn1* appressoria at 16 h, compared to 86.4% of Guy11 appressoria (S6D and S6F Fig). These results indicated that MoCrn1 is indispensable for an efficient degradation of glycogen and lipid necessary for the appressorial turgor generation.

We further evaluated the Δ*Mocrn1* mutant for pathogenicity on rice. The conidial suspensions from Guy11, Δ*Mocrn1,* and the complemented strain were sprayed onto the susceptible rice cultivar CO-39. Δ*Mocrn1* produced fewer lesions than Guy11 and the complemented strain, which were confirmed by lesion quantification (Fig 7A). We also performed rice sheath penetration assays by observing 100 appressoria each strain and classifying invasive hyphae (IH) types as previously described [13]. We observed that over 40% of Δ*Mocrn1* appressoria were defective in penetration and 55.6% of appressoria that penetrated and formed less extended IH. In contrast, 90% of Guy11 appressoria successfully penetrated rice cells and about 50% of that produced strong IH (Fig 7B).

**Fig 7.**
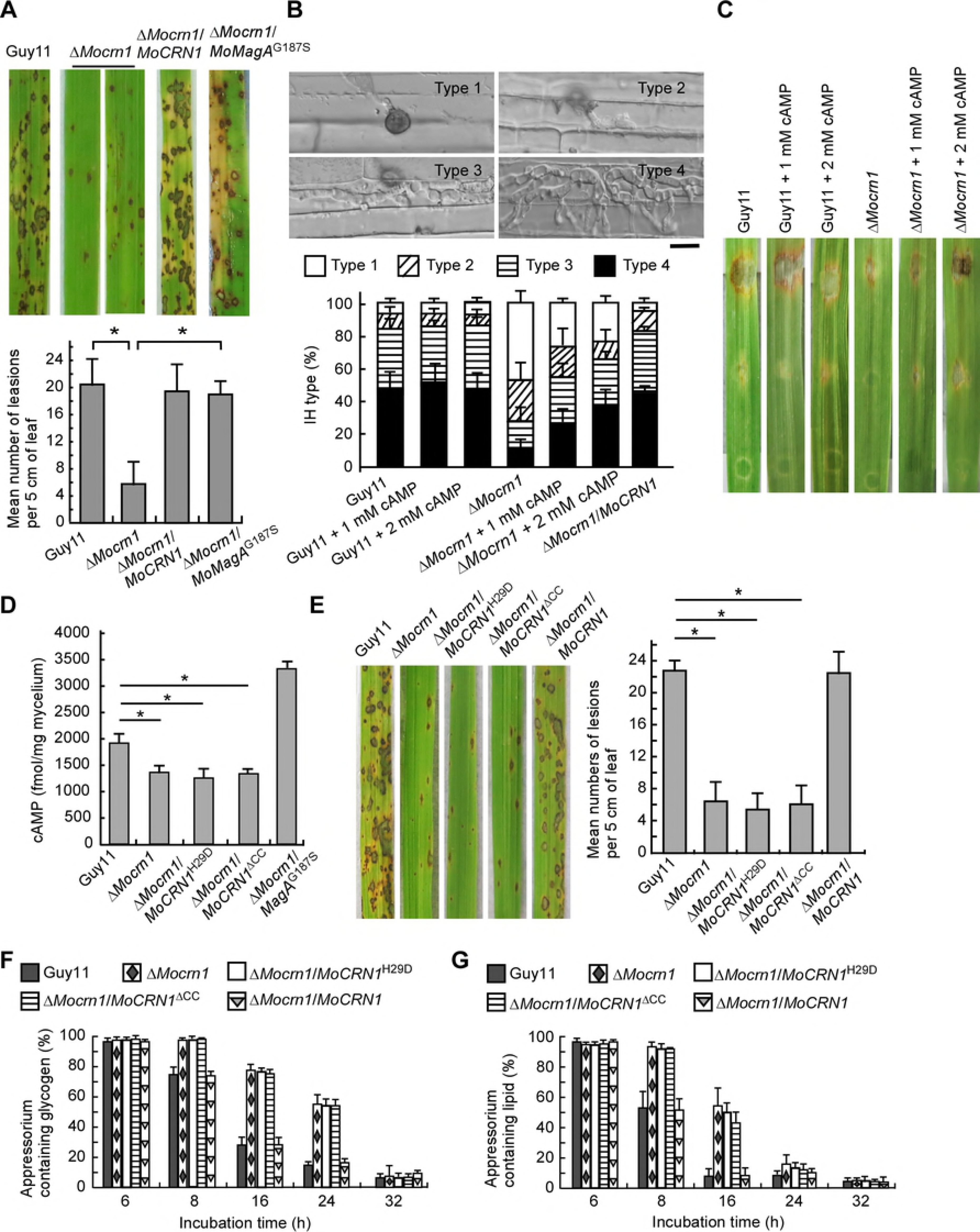
MoCrn1 contributes to pathogenicity through its function in regulating the cAMP level. **(A)** Pathogenicity assay was conducted by spraying conidial suspensions (5×10^4^ conidia/ml) onto two-week old rice seedlings (CO-39). Mean number of lesions per 5 cm length of leaf were quantified. Expressing MoMagAG187S in Δ*Mocrn1* suppressed the defects in infection. **(B)** Penetration assays with rice sheath tissues. Micrographs show the percentages of 4 types of IH observed at 36 hpi. Bars = 10 μm. 1 or 2 mM 8-Br-cAMP addition could promote penetration for Δ*Mocrn1*. **(C)** Pathogenicity assay was conducted with detached barley leaves. Addition of 1 mM and 2 mM 8-Br-cAMP enhanced Δ*Mocrn1* infection. **(D)** Bar chart shows the intracellular cAMP levels in mycelium of the strains. **(E)** Mutation of H29D and CC deletion for MoCrn1 caused defects in pathogenicity. Quantification for the lesions per 5 cm length, leaf is given. **(F and G)** Bar charts show the percentages of appressoria containing glycogen and lipids at different time points. The Δ*Mocrn1* mutant, Δ*Mocrn1*/*MoCRN1*^H29D^ and Δ*Mocrn1*/*MoCRN1*^ΔCC^ strains delayed the degradation of glycogen and lipids.

### MoCrn1 is involved in appressorium development and pathogenicity through regulating intracellular cAMP levels

To explore whether MoCrn1 regulates turgor generation involving the process of cAMP signaling, the incipient collapse assay was performed. We found that exogenous 8-Br-cAMP could suppress the defect of Δ*Mocrn1* in turgor generation (S6B Fig). The numbers of the collapsed appressoria in the Δ*Mocrn1* mutant were reduced by 20% and 10% with 1 and 2 mM cAMP, respectively, compared to those without 8-Br-cAMP. In addition, the Δ*Mocrn1* mutant appressorium underwent successful glycogen and lipid breakdown following 8 and 16 h, respectively, following treatment with 5 mM 8-Br-cAMP (S6E and S6F Fig). Furthermore, 1 or 2 mM 8-Br-cAMP addition to the conidia suspensions in the inoculation of detached barley leaves could suppress the defect of Δ*Mocrn1* in infection to some degree (Fig 7C). This result was also confirmed by the penetration assay, in which 8-Br-cAMP treatment restored the penetration defect to almost 80% of the Δ*Mocrn1* appressoria in comparison to 43 ± 4.9% of Δ*Mocrn1* without cAMP (Fig 7B). This is similar to the effect of the Δ*Mocrn1* mutant that expresses the constitutively activated allele of MoMagA, MoMagA^G187S^ (S6G and S6H Fig and Fig 7A).

### MoCrn1 function is dependent on its protein binding ability

To examine the ability of MoCrn1 in binding multiple proteins, we identified putative actin binding domains and characterized their function. Human coronin Arg^29^ and Arg^30^ are thought to be important for the interaction with F-actin [27, 28]. The alignment showed that a majority of coronins contain a conserved basic amino acid at these two positions (Fig 8A). In addition, the C-terminal coiled-coil (CC) domain is important for coronins to interact with the actin nucleation complex Arp2/3 [29]. Accordingly, we mutated His^29^ to Asp^29^ and deleted the CC domain of MoCrn1, and fused the mutant proteins with GFP (Fig 8B). We found that MoCrn1^H29D^ and MoCrn1^ΔCC^ mutants had completely altered actin-like localization patterns (Fig 8C). To further analyze the effects of these mutant alleles, we performed the co-IP assay and found that MoCrn1^H29D^ and MoCrn1^ΔCC^ mutants failed to interact with MoRgs7, MoMagA, and MoCap1 (Fig 8D 8E, and 8F).

**Fig 8.**
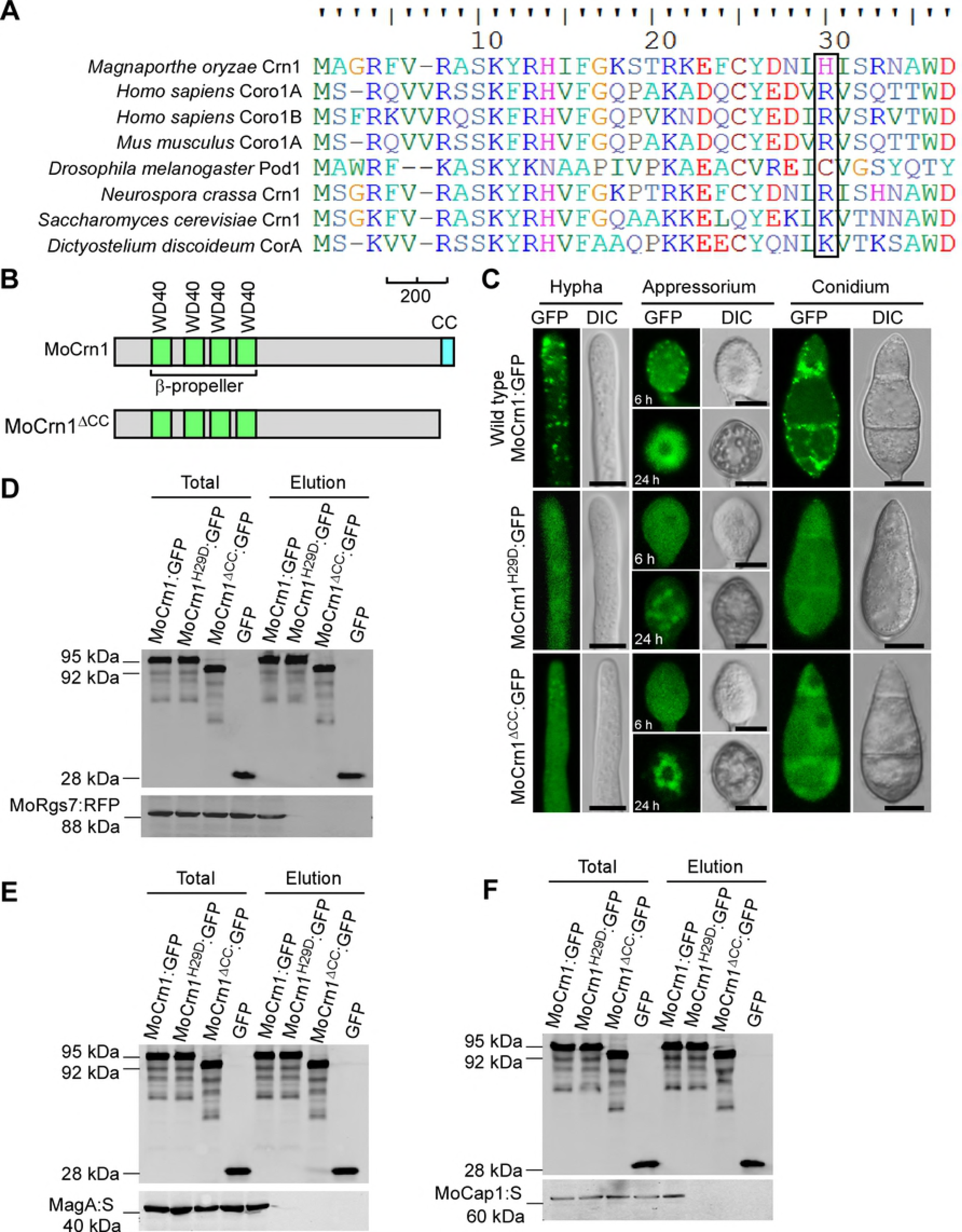
Dissection of MoCrn1 residues and domains crucial for protein-protein interactions. **(A)** Comparison of the actin binding position of MoCrn1 (the 29^th^ His) to that of coronins in other species. Accession number: *Homo sapiens* Coro1A (NP_009005.1) and Coro1B (NP_065174.1), *Mus musculus* Coro1A (:NP_034028.1), *Drosophila melanogaster* Pod1 (NP_001245554.1), *Neurospora crassa* Crn1 (XP_956587.2), *Saccharomyces cerevisiae* Crn1 (NP_013533.1) and *Dictyostelium discoideurm* CorA (CAA43707.1). **(B)** Schematic representation of MoCrn1 and MoCrn1^ΔCC^ draw by using the following colors: green, WD40; blue, coiled-coil. Scale bar, 200 amino acids. **(C)** Images show the localization patterns of MoCrn1:GFP, MoCrn1^H29D^:GFP and MoCrn1^ΔCC^:GFP in hypha, conidium and appressorium. **(D, E and F)** Co-IP assays for assaying the interaction of MoCrn1 (tagged with GFP), MoCrn1^H29D^ (tagged with GFP) and MoCrn1^ΔCC^ (tagged with GFP) with MoRgs7 (tagged with RFP), MoMagA (tagged with S) and MoCap1 (tagged with S).

We also expressed MoCrn1^H29D^ and MoCrn1^ΔCC^ mutants in Δ*Mocrn1*. FRAP analysis showed that the expression of MoCrn1^H29D^ and MoCrn1^ΔCC^ caused no effect on delayed endocytosis of MoRgs7 and MoMagA in Δ*Mocrn1* (Fig 9). HPLC analysis revealed cAMP levels of the strain expressing MoCrn1^H29D^ or MoCrn1^ΔCC^ comparable to that of the Δ*Mocrn1* mutant (Fig 7D). Moreover, virulence and the degradation of appressorial glycogen and lipid in the MoCrn1^H29D^ and MoCrn1^ΔCC^ strains were also indistinguishable from those of the Δ*Mocrn1* mutant (Fig 7E 7F, and 7G). Taken together, these results suggested that MoCrn1 function is dependent on its ability to interact with F-actin, MoRgs7, MoMagA, and MoCap1.

**Fig 9.**
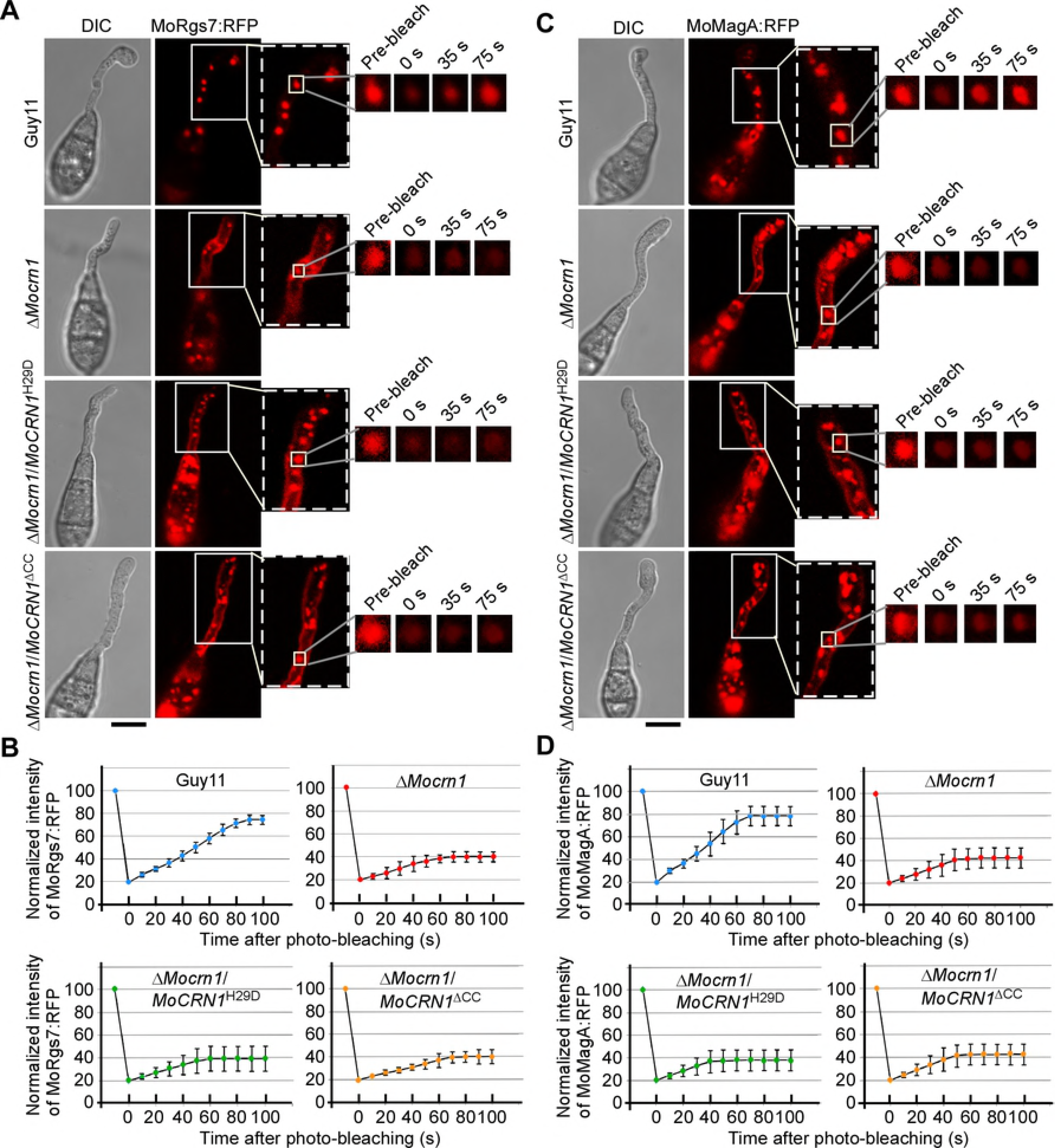
The 29^th^ His and CC domain are important for MoCrn1 function in MoRgs7 and MoMagA endocytosi. **(A)** FRAP for measuring the recovery of MoRgs7:RFP fluorescence on endosomes of germ tubes. FRAP analysis was conducted at 3 h post-germination. The representative images of FRAP were shown and the selected areas were measured for fluorescence recovery after photobleaching. Bar = 5 μm. **(B)** The normalized FRAP curves of MoRgs7:RFP were fitted with measuring 15 regions from different cells. Bars = 5 μm. **(C)** FRAP for measuring the recovery of MoMagA:RFP fluorescence on endosomes of germ tubes. FRAP analysis was conducted at 3 h post-germination. The representative images of FRAP were shown and the selected areas were measured for fluorescence recovery after photobleaching. Bar = 5 μm. **(D)** The normalized FRAP curves of MoMagA:RFP were fitted with measuring 15 regions from different cells. Bars = 5 μm.

## Discussion

We here investigate the distinct functional mechanism of 7-TM-containing protein MoRgs7 beyond its RGS functions. We found that MoRgs7 has a GPCR-like endocytosis pattern and is predominantly localized to late endosomes similar to other signaling proteins including MoRgs1, MoMagA, and MoMac1. Such late endosome localizations of signaling proteins are critical to GPCR function and for cAMP signaling transduction. Our results further showed that MoRgs7 couples with MoMagA to undergo endocytosis. Interestingly, by inhibiting endocytosis, we could observe increased PM localization of MoRgs7 and MoMagA. And by inhibiting trafficking from the early endosomes to the late endosomes, we could observe the early endosome localization of MoRgs7 and MoMagA.

Understanding how pathogen receptors recognize the plant surface signal has beneficial effect on controlling rice disease at early stages. Our results provide evidences that MoRgs7 serves as a GPCR-like receptor to detect environmental hydrophobic cues. The affinity precipitation assay with phenyl-agarose gel beads indicates that MoRgs7 has strong ability to form hydrophobic interaction with hydrophobic materials, revealing that MoRgs7 can form interaction with hydrophobic surface when MoRgs7 is localized to PM. Importantly, disruption of such hydrophobic interaction during *M. oryzae* germinating on hydrophobic surface led to the aberrant appressorium formation. We also noted that the Δ*Morgs7* mutant developed defective appressoria, even though no decrease in appressorium formation frequency. Based on these studies, we concluded that forming hydrophobic interactions with hydrophobic surface by MoRgs7 and other membrane proteins is a critical step in recognizing hydrophobic surface cues.

We reasoned that MoRgs7 may undergo a process similar to mammalian GPCRs. In mammalian cells, a ligand binding to a GPCR can activate it by inducing conformational changes in GPCR. The active GPCRs can activate the Gα proteins by exchanging GDP of Gα proteins to GTP Gα. Meanwhile active GPCRs are transported by endocytosis to sustain downstream signaling, recycling, or be degraded from endosomes [30]. Considering our studies and findings in mammalian cells, we proposed a functional model of MoRgs7 (Fig 10) in which MoRgs7 acts as a GPCR during appressorium development to interact with the hydrophobic surface. Subsequently, this interaction induces MoRgs7-MoMagA endocytosis that is recruited by MoCrn1. Thereby, MoRgs7 facilitates activating cAMP signaling from endosomes along with MoMagA. Conversely, MoRgs7 may elevate its GAP activity when cAMP signaling is fully activated. Thus, MoRgs7 has dual roles in regulating signal transduction. However, how MoRgs8 that also contains 7-TM domain but lacks sensory functions is not understood. MoRgs8 was distributed in the cytoplasm of germ tubes but did not undergo endocytosis. We speculated that MoRgs8 could be involved in a mechanism that differs from MoRgs7. For example, MoRgs7 does not respond to surface hydrophilic signals, could this be the role of MoRgs8?

**Fig 10.**
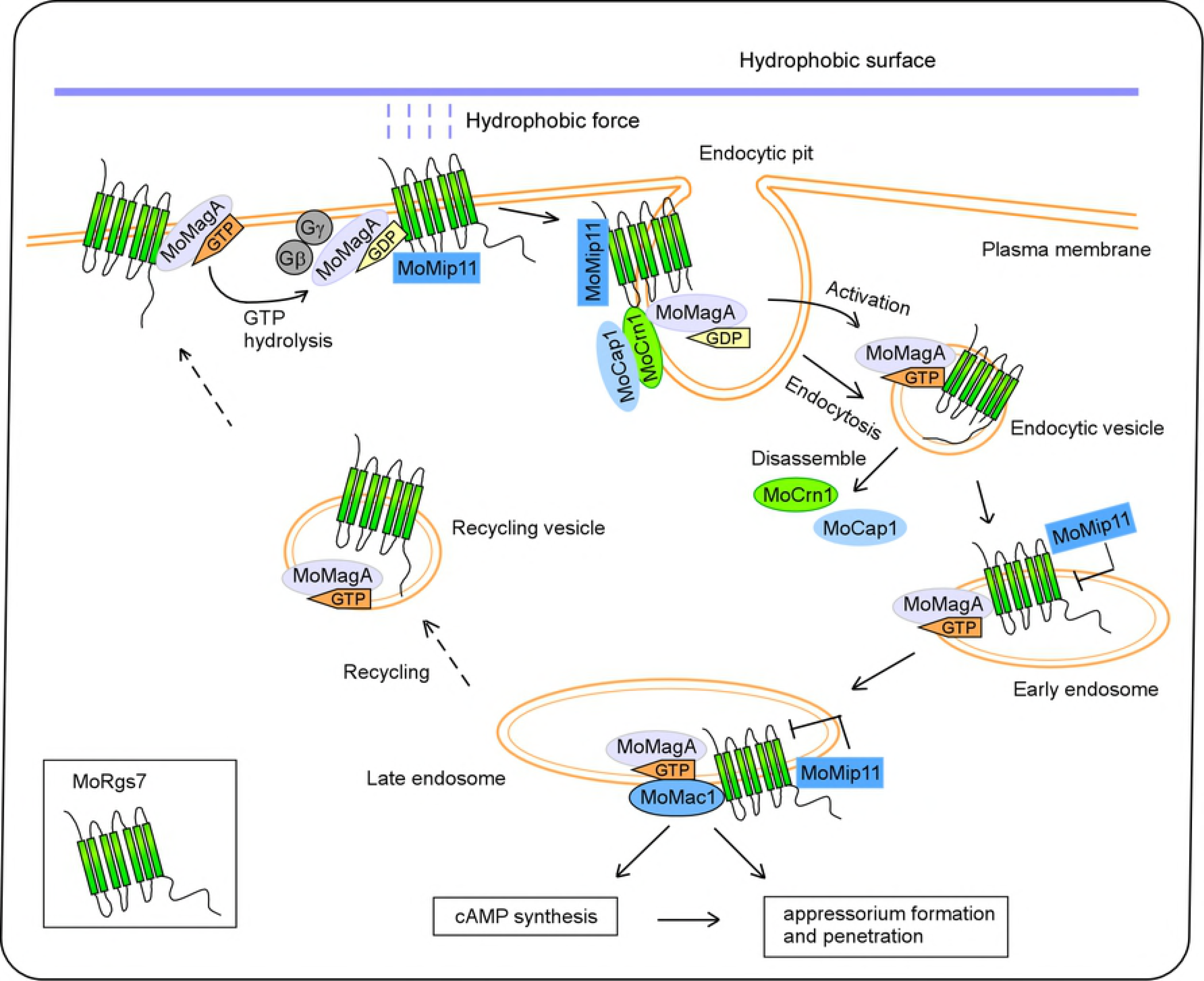
Model of MoRgs7 mediating perception of hydrophobic surface and promoting cAMP signaling by transport through the endocytic pathway. When MoRgs7 forms hydrophobic interaction with hydrophobic surface, MoCrn1 interacts with MoRgs7, MoMagA and MoCap1 and directs them to endocytic sites. Subsequently MoRgs7 couples with MoMagA to undergo endocytosis to late endosomes. As MoMagA dissociates from Gβ and Gγ subunits and becomes GTP bound MoMagA, MoRgs7 and MoMagA function to promote MoMac1 activation and cAMP signaling, which led to the proper appressorium development and pathogenicity. To sustain signal transduction, MoMip11 that interacts to MoRgs7 can inhibit MoRgs7 interacting with GTP bound MoMagA to prevent GTP hydrolysis in MoMagA. As MoRgs7 and MoMagA are sent back to the plasma membrane via recycling pathways, MoMagA binds to Gβ-Gγ again following GTP hydrolysis in MoMagA.

There was precedence that endocytosis of RGS proteins plays a role in promoting Gα-mediated signaling. In *Arabidopsis thaliana*, in response to glucose RGS protein AtRgs1 internalizes via endocytosis to uncouple itself from Gα protein AtGPA1 anchored in the PM, leading to AtGPA1 sustain activation. And this process is required for both G-protein-mediated sugar signaling and cell proliferation [31]. However, other details including the initiation of MoRgs7-MoMagA complex disassembly following endocytosis remain not understood. We recently reported a distinct mechanism of how *M. oryzae* might negatively regulate the GAP activity of MoRgs7. This mechanism implicates the MoMip11 protein that interacts with MoRgs7 and the GDP bound MoMagA, but not the GTP bound MoMagA (Fig 10) [10]. MoMip11 prevents MoRgs7 from interacting with the GTP bound MoMagA, therefore interfering with MoRgs7 GAP function by sustaining MoMagA activation [10].

To further investigate the physiological function of MoRgs7 and MoMagA endocytosis, we identified and characterized coronin protein MoCrn1 and found that it regulates MoRgs7 and MoMagA endocytosis. MoCrn1 is localized to actin patches that represent endocytic sites, interacting with MoRgs7 and MoMagA. Disruption of MoCrn1 by gene deletion or point mutations (H29D mutation and CC domain deletion) not only attenuated MoRgs7 and MoMagA endocytosis, but also led to a decreased cAMP level that is lower than the threshold for proper appressorium development. Our results support that MoRgs7 and MoMagA endocytosis regulated by MoCrn1 facilitates initiating cAMP signaling and appressorium development.

Coronin proteins are known as regulators of the cytoskeleton and membrane trafficking in a number of species including yeast, *Neurospora crassa*, *Dictyostelium discoideum*, *Drosophila*, and human [19, 32]. In *D. discoideum* and mammalian cells, coronins have evolved to be modulators of signal transduction. Those coronins are critical for Rac1 GTPase activation and Rac1-dependent signaling [28, 33]. Additionally, upon cell surface stimulation coronin 1 interacts with and activates Gα to stimulate cAMP/PKA pathway in neuronal cell, even though how coronin 1 activates Gα is less clear [34]. Compared to those studies, our work revealed that MoCrn1 is involved in a distinct mechanism to facilitate Gα-cAMP signaling. MoCrn1 has an adaptor protein-like function by directing MoRgs7 and MoMagA to endocytic pits to promote their internalization, this function thereby allows MoCrn1 to have a role in facilitating cAMP signaling. Interestingly, MoCrn1 also interacts with MoCap1 that is thought as one of activators of MoMac1 [6]. Based on the above, we proposed that MoCrn1 is likely to be a hub or organizing protein of the network of MoRgs7-MoMagA-MoCap1.

## Materials and Methods

### Strains and culture conditions

The *M. oryzae* Guy11 strain was used as wild type for transformation in this study. For vegetative growth, small agar blocks were taken from the edge of 7-day-old cultures and cultured in liquid CM medium for 48 h. For conidiation, strains were cultured on SDC plates at 28°C for 7 days in the dark, followed by constant illumination for 3 days [8, 13, 26, 35–38].

### Targeted *MoCRN1* deletion and the Δ*Mocrn1* mutant complementation

The *MoCRN1* deletion mutant was generated using the standard one-step gene replacement strategy [39]. First, two approximate 1.0 kb of sequences flanking of *MoCRN1* (MGG_06389) were amplified with two primer pairs *MoCRN1*-F1/*MoCRN1*-R1, *MoCRN1*-F2/*MoCRN1*-R2, the resulting PCR products ligated with the *HPH* cassette released from pCX62. The protoplasts of wild type Guy11 were transformed with the vectors for targeted gene deletion by inserting the hygromycin resistance *HPH* marker gene cassette into the two flanking sequences of the *MoCRN1* gene. For selecting hygromycin-resistant transformants, CM plates were supplemented with 250 μg/ml hygromycin B (Roche, USA).

To generate complementary construct pYF11-*MoCRN1*, the gene sequence containing the *MoCRN1* gene and 1.0 kb native promoter was amplified with *MoCRN1*-comF/ *MoCRN1*-comR. Yeast strain XK1-25 was co-transformed with this sequence and *Xho*I-digested pYF11 plasmid. Then the resulting yeast plasmid was expressed in *E. coli*. To generate the complementary strain, the pYF11-*MoCRN1* construct was introduced into the Δ*Mocrn1* mutant and pYF11 contains the bleomycin-resistant gene for *M. oryzae* transformants screen [26, 39].

### Southern blot analysis

*Eco*RV was used to digest the genomic DNA from Guy11 and the Δ*Mocrn1* mutant. The digest products were separated in 0.8% agar gel and were hybridized with the *MoCRN1* gene probe. The probe was designed according to the disruption strategy and was amplified from Guy11 genomic DNA using primers *MoCRN1*-InterF/*MoCRN1*-InterR. To confirm *MoCRN1* replacements, labeled *MoCRN1* probe was used to hybridize the *Eco*RV-digested genomic DNA from the Δ*Mocrn1* mutant and wild-type Guy11. The copy number of the *HPH* gene in the Δ*Mocrn1* mutant was detected using labeled *HPH* fragments that amplified from the plasmid of pCB1003 with primers FL1111/FL1112. The whole hybridization was carried out according to the manufacturer’s instruction for DIG-High Prime.

### Pathogenicity assay

The conidia were suspended in a 0.2% (w/v) gelatin solution (5×10^4^ spores/ml), then the solutions were sprayed onto 2-week-old seedling of susceptible rice (*Oryza sativa* cv. CO-39) and also inoculated into 3-week-old rice CO-39 as described. Then the plants were incubated at 25°C with 90% humidity in the dark for the first 24 h, followed by a 12 h/12 h light/dark cycle. Lesions were observed after 7 days of incubation [36]. For pathogenicity assay with detached barley leaves [35], three 20 μl droplets of the conidia suspensions (1×10^5^, 1×10^4^, 1×10^3^ spores/ml, respectively) added cAMP solution or not, were placed onto the upper side of the 7-day-old barley (cv. Four-arris) leaves. Then the leaves were incubated at 25°C with 90% humidity and in the dark for the first 24 h, followed by a 12 h/12 h light/dark cycle. Lesions were observed after 5 days of incubation.

### Glycogen and lipid staining during appressorium development

To visualize glycogen, the samples were stained by iodine solution containing 60 mg/ml KI and 10 mg/ml I_2_ for 1 min. Nile red solution consisting of 50 mM Tris/maleate buffer (pH 7.5) and 2.5 mg/ml Nile red (9-diethylamino-5H-benzo-a-phenoxazine-5-one, Sigma), was used to treat the samples for 3 min, then the samples were examined under a fluorescence microscope with RFP channel [13, 21, 25].

### Co-IP assay

The DNA fragments for expressing GFP fusion proteins were respectively inserted into the pYF11 construct that contains bleomycin resistant gene and G418 resistance gene, and the DNA fragments for expressing S-tag fusion proteins were respectively inserted into the pXY203 construct hat contains hygromycin gene. Then the constructs for expressing GFP and S-tag fusion proteins were co-transformed into wild-type strain Guy11, and the transformants resistant to hygromycin and bleomycin or G418 were isolated. The total protein of the transformants was extracted from mycelium using protein lysis buffer [1 M Tris-Cl (pH7.4), 1 M NaCl, 0.5 M EDTA, 1% Triton×100] and incubated with anti-GFP agarose beads (GFP-Trap, Chromotek, Martinsried, Germany) for 4 h, followed by washing beads with washing buffer (50 mM Tris HCl, 150 mM NaCl, pH 7.4) for 4 times. The proteins that bind to the beads were eluted by 0.1 M glycine HCl (pH 3.5) and were probed by anti-GFP and anti-S antibodies.

### Binding assay for MoCrn1 and MoAct1 interaction

*MoCRN1* and *MoACT1* full-length cDNAs were cloned and inserted into pGEX4T-2 and pET32a, respectively. These constructs were transformed into *E. coli* strain BL21 for expressing proteins. Bacterial lysate containing GST:MoCrn1 protein was incubated with 30 μl GST agarose beads for 2 h. Then the beads were washed by washing buffer for 4 times and incubated with His:MoAct1 protein for 2 h, followed by washing beads with using washing buffer (50 mM Tris HCl, 150 mM NaCl, pH7.4) for 4 times again. The beads were boiled to elute proteins, and eluted proteins (output) were probed with anti-GST and anti-His antibodies.

### Yeast two-hybrid assay

Constructs of BD:*MoMagA,* BD:*MoMagB* and BD:*MoMagC* were used in previous experiments and kept in our lab. Full-length cDNAs of *MoCRN1* was cloned and inserted into pGADT7 (AD) vector. Full-length cDNAs of *MoCAP1*, *MoMagA*^G187S^, *MoMagA*^Q208L^ and *MoACT1* genes were inserted into pGBKT7 (BD) vector. To examine the interaction of proteins, the AD and BD constructs were co-transformed into yeast strain AH109 and the transformants were grown on SD-Trp-Leu medium. Then the Trp+ and Leu+ transformants were isolated and assayed for growth on SD-Trp-Leu-His-Ade medium added X-α-Gal.

### FRAP analysis

The germinated conidia with 3 h of incubation on hydrophobic or hydrophilic surfaces were treated with cycloheximide and benomyl as described. FRAP were performed using a fluorescence microscope Zeiss LSM710. Regions containing MoRgs7:RFP and MoMagA:RFP in germ tube were selected for photo-bleaching. Photobleaching was carried out using an Argon-multiline laser at a wavelength of 561 nm with 80% laser power and 80 iterations in ROI. Images were acquired with 2% laser power at a wavelength of 555 nm every 5 sec. For quantitative analyses, fluorescence intensity was measured using the ZEISS ZEN blue software and fluorescence recovery curves were fitted using following formula: F(t) = F_min_ + (F_max_ – F_min_)(1−exp^−kt^), where F(t) is the intensity of fluorescence at time t, F_min_ is the intensity of fluorescence immediately post-bleaching, F_max_ is the intensity of fluorescence following complete recovery, and k is the rate constant of the exponential recovery [40]. Mobile Fraction was calculated as the following formula: Mf = (F_end_ – F_0_)/(F_pre_ – F_0_), where F_end_ is the stable fluorescent intensity of the punctae after sufficient recovery, F_0_ is the fluorescent intensity immediately after bleaching, and F_pre_ is the fluorescent intensity before bleaching [41].

### Assays with drugs or inhibitors

Latrunculin B (Cayman, USA) is dissolved in DMSO at a concentration of 25 mg/ml. Conidia incubated on the coverslips with hydrophobic surface were treated with LatB (final concentration 0.1 μg/ml) for 30 min, while the controls were treated with 5% DMSO. Then samples were washed with distilled water. Cycloheximide (MedChemExpress, USA) was solved in distilled water and the germinated conidia were treated with a final concentration 10 μg/ml for 10 min. Then samples were washed with distilled water. Benomyl (Aladdin, Shanghai, China) was solved in 0.1% DMSO and added to germinated conidia with a final concentration 1μg/ml. Then the samples were washed with distilled water. EGA (Merck, USA) was solved in 5% DMSO and was applied to samples with concentration 5 μg/ml for 1 h.

### Affinity precipitation of MoRgs7:GFP with Phenyl-agarose gel beads

The total proteins were extracted from the Guy11 strain expressing MoRgs7:GFP or GFP, respectively, and were incubated with 100 mg of Phenyl-agarose beads (Senhui Microsphere Tech, Suzhou, China) in 1.5 ml microcentrifuge tubes at 10°C for 16 h. After incubation, the tubes were centrifuged (13000 g, 5 min) to remove the suspension. The beads were then gently washed with a series of aqueous solutions with different concentrations of NaCl and MgSO4 (1.5/1.0/0.8/0.5/0.3/0.2/0.1 M NaCl and MgSO4, 10 mM HEPES, pH 7.0), respectively, for 3 times to remove the unbound proteins. 100 μl of 1% SDS solution was added to the washed beads, followed by boiling the SDS solution and beads for 10 min to obtain elution, which was examined by western-blot using anti-GFP antibody.

### Assays with fluorescence microscope and calculation of Pearson correlation coefficient for co-localization

All the samples were observed under a confocal fluorescence microscope (Zeiss LSM710, 63× oil). The filter cube sets: GFP (excitation spectra: 488 nm, emission spectra: 510 nm), RFP (excitation spectra: 555 nm, emission spectra: 584 nm). Exposure time: 800 ms. ImageJ software was applied to calculate Pearson correlation coefficient for analyzing colocalization of GFP fusion protein with RFP fusion protein. One area of interest was photographed with GFP and RFP channels respectively and photographs were opened using ImageJ software. Picture type was set at 8 bits. The “colocalization finder” in “plugin” section was applied to the pictures and Pearson correlation coefficient was calculated.

### cAMP extraction and high-performance liquid chromatography (HPLC) analysis

All of the strains were cultured on CM medium at 28°C, were cut into 1×1 mm squares, and were cultured in liquid CM for another 2 days. Filtering to collect mycelium and quickly ground into powder in liquid N_2_. 1 mg of mycelium was mixed with 20 μl of 6% TCA solution. Samples were centrifuged (1,377 × g, 15 min), the top layers were collected and were washed twice with five times the volume of anhydrous ether. The pellet was collected for HPLC. HPLC analysis was done with a programmable Agilent Technology Zorbax 1200 series liquid chromatograph. The solvent system consisted of methanol (90%) and water (10%), at a flow rate of 1 ml per minute; 0.1 mg of cAMP solution per milliliter was eluted through the column (SBC18, 5 μl, 4.6 × 250 mm) and was detected at 259 nm UV. Each sample was eluted through the column in turn and peak values were detected with the same time as the standard [42].

### Construction of vectors used to express fluorescent proteins

For construction of pHZ65:*MoMagA* vector used to express MoMagA-N’YFP, the N’YFP sequence was inserted into the alphaB-alphaC loop of MoMagA as described [43], then the MoMagA sequence containing N’YFP and the native promoter was fused with the pYF11 plasmid. For construction of vector used to express MoMagA:RFP, the RFP sequence was also inserted into the alphaB-alphaC loop of MoMagA. Then the MoMagA sequence containing the native promoter was fused with pYF11 plasmid. For construction of other vectors used to express proteins tagged with RFP or GFP, RFP or GFP was fused to protein sequence C-terminals, then protein sequences containing their native promoters were fused with the pYF11 plasmid.

### GenBank accession number

*MoRGS7* (MGG_11693), *MoRGS8* (MGG_13926), *MoMagA* (MGG_01818), *MoCRN1* (MGG_06389), *MoCAP1* (MGG_01722)

## Acknowledgments

We thank Dr. Naweed I Naqvi of the National University of Singapore for providing GFP:MoRab5 and GFP:MoRab7 plasmids.

## Supporting information

**S1 Fig. MoRgs7 and MoRgs8 are predicted to contain a 7 transmembrane domain. (A)** The analysis results to confirm the 7-TM domain in MoRgs7 were yielded by the websites http://mendel.imp.univie.ac.at/sat/DAS/DAS.html and http://www.cbs.dtu.dk/services/TMHMM. **(B)** The analysis results to confirm the 7-TM domain in MoRgs8.

**S2 Fig. The RGS and 7-TM domain is required for MoRgs7 function. (A)** The schematic representations of MoRgs7, 7-TM and MoRgs7^Δ7-TM^ were drawn with green that represents 7-TM and blue that represents RGS domain. **(B)** Bar chart shows the intracellular cAMP levels in the mycelium. The values were recorded from three independent experiments. NS represents no significant differences. **(C)** Bar chart shows the percentages of the conidia generating two appressoria. The values were recorded from three independent experiments. NS represents no significant differences. **(D)** Pathogenicity assay was conducted by spaying conidial suspensions (5×10^4^ conidia/ml) onto two-week old rice seedlings (CO-39). **(E)** Mean number of lesions per 5 cm length of leaves were quantified for **(D)**.

**S3 Fig. MoRgs7 and MoMagA are independent of each other in internalization.(A)** The representative images of FRAP analysis for MoRgs7:RFP were shown and the selected areas were measured for fluorescence recovery after photobleaching. FRAP analysis was conducted at 3 h post-germination. Bar = 5 μm. **(B)** The normalized FRAP curve of MoRgs7:RFP were fitted with measuring 15 regions from different cells. **(C)** The representative images of FRAP analysis for MoMagA:RFP were shown and FRAP analysis was conducted at 3 h post-germination. Bar = 5 μm. **(D)** The normalized FRAP curve of MoMagA:RFP were fitted with measuring 15 regions from different cells.

**S4 Fig. Targeted *MoCRN1* deletion was confirmed by Southern blot analysis.** Southern blot analysis of the *MoCRN1* gene deletion mutants with gene specific probe (probe1) and hygromycin phosphotransferase (HPH) probe (probe2). Thick arrows indicate the orientations of the *MoCRN1* and HPH genes. Thin lines below the arrows indicate sequence-specific gene probes.

**S5 Fig. MoCrn1 is co-localized with F-actin and has actin and microtubule-associated functions. (A)** MoCrn1 is co-localized with F-actin in appressorium, hypha and conidium. Scale bar for the appressorium, 10 μm. Bar of hypha, 5 μm. Bar of conidium, 5 μm. **(B)** The yeast two-hybrid assay for examining the interaction of MoCrn1 with actin protein MoAct1. The yeast transformants were isolated from SD-Leu-Trp plates, following growing on SD-Leu-Trp-His-Ade plates containing X-α-Gal for examining β-galactosidase activity. **(C)** Binding assay for examining the interaction of MoCrn1 with actin protein MoAct1. Input represents the proteins extracted from the *E. coli* BL21 strains expressing GST-MoCrn1 or His-MoAct1. Output represents the proteins eluted from the GST-beads used to bind GST-MoCrn1. Those proteins were probed by using GST-antibody and His-antibody. **(D)** Images show actin structures labeled by lifeact:RFP in appressoria. Bar = 5 μm. **(E)** Images show actin structures labeled by lifeact:RFP in conidia. Bar = 5 μm. **(F)** Images show actin structures labeled by lifeact:RFP in hyphae. Bar = 5 μm. **(G)** The assay for determining sensitivity to benomyl. The yeast wild-type BY4741, the Δ*crn1* mutant and the Δ*crn1*Δ/*MoCRN1* strains were grown on SD plates containing 0, 10, 20 and 30 μg/ml benomyl for 3 days. **(H)** The colonies of Guy11, the Δ*Mocrn1* mutant and the complemented strain grew on CM plates containing 0, 0.6, 1.0 and 1.2 μg/ml benomyl for 7 days. Bar chart shows the inhibition rate. The experiment was repeated three times.

**S6 Fig. MoCrn1 affects glycogen and lipid degradation during appressorium development through regulating cAMP synthesis. (A)** Bar chart shows the intracellular cAMP levels in mycelium of Guy11, the Δ*Mocrn1* mutant and the complemented strain. Asterisks represent significant differences (P < 0.01). **(B)** Incipient collapse assay was conducted with 1, 2 and 3 M glycerol solution to examine the appressorial turgor level. Bar chart shows the percentages of collapse appressoria upon glycerol solution treatment and 8-Br-cAMP addition decreased the collapse rate of Δ*Mocrn1* appressoria. 200 appressoria were observed for each sample and the experiment was repeated three times. **(C)** Micrographs show the glycogen distribution in Guy11 and Δ*Mocrn1* at different time points. The conidia of Guy11 and Δ*Mocrn1* were allowed to germinate on hydrophobic surface, and glycogen could be visualized by iodine solution staining. Bar = 10 μm. **(D)** Micrographs show the lipid distribution in Guy11 and Δ*Mocrn1* at different time points. Lipid bodies were visualized by Nile red staining. Bar = 10 μm. **(E and F)** Bar charts show the percentages of appressoria containing glycogen and lipids at different time points. The 5 mM cAMP treatment significantly promoted degradation of glycogen and lipids in Δ*Mocrn1* appressoria. 200 appressoria were observed for each sample and the experiment was repeated three times. **(G and H)** Bar charts show the percentages of appressoria containing glycogen and lipids at different time points. Expressing MoMagA^G187S^ in Δ*Mocrn1* significantly promoted degradation of glycogen and lipid in appressoria. 200 appressoria were observed for each sample and the experiment was repeated three times.

**S7 Fig. MoCrn1 interacts with MoCap1 and controls MoCap1 localization.** The localization pattern of MoCap1 was severely disrupted in mature appressorium, conidium and hypha of Δ*Mocrn1*. Bars = 5 μm.

**S8 Fig. Δ*Mocrn1* was slightly delayed in appressorium formation.** Appressoria that formed in hydrophobic surfaces were observed at 4, 5 and 6 h post-germination. Bar = 15 μm. The formation percentages were quantified by observing 200 appressoria for each sample and the experiment was repeated three times.

**S1 Table. Primers used in this study.**

